# Transcription inhibitors with XRE DNA-binding and cupin signal-sensing domains drive metabolic diversification in *Pseudomonas*

**DOI:** 10.1101/2020.07.29.226225

**Authors:** Julian Trouillon, Michel Ragno, Victor Simon, Ina Attrée, Sylvie Elsen

## Abstract

Transcription factors (TFs) are instrumental in the bacterial response to new environmental conditions. They can act as direct signal sensors and subsequently induce changes in gene expression leading to physiological adaptation. Here, by combining RNA-seq and DAP-seq, we studied a family of eight TFs in *Pseudomonas aeruginosa*. This family, encompassing TFs with XRE-like DNA-binding and cupin signal-sensing domains, includes the metabolic regulators ErfA, PsdR and PauR and five so far unstudied TFs. The genome-wide delineation of their regulons identified 39 regulatory interactions with genes mostly involved in metabolism. We found that the XRE-cupin TFs are inhibitors of their neighboring genes, forming local, functional units encoding proteins with functions in condition-specific metabolic pathways. The phylogenetic analysis of this family of regulators across the *Pseudomonas* genus revealed a wide diversity of such metabolic regulatory modules and identified species with potentially higher metabolic versatility. Numerous uncharacterized XRE-cupin TFs were found near metabolism-related genes, illustrating the need of further systematic characterization of transcriptional regulatory networks in order to better understand the mechanisms of bacterial adaptation to new environments.

**IMPORTANCE:** Bacteria of the *Pseudomonas* genus, including the major human pathogen *P. aeruginosa*, are known for their complex regulatory networks and high number of transcription factors, which contribute to their impressive adaptive ability. However, even in the most studied species, most of the regulators are still uncharacterized. With the recent advances in high-throughput sequencing methods, it is now possible to fill this knowledge gap and help understanding how bacteria adapt and thrive in new environments. By leveraging these methods, we provide an example of a comprehensive analysis of an entire family of transcription factors and bring new insights into metabolic and regulatory adaptation in the *Pseudomonas* genus.

## INTRODUCTION

The regulation of gene transcription is a key mechanism in the evolution and adaptation of bacteria to a wide array of environments. Transcriptional regulation involves the interaction between *trans*-acting regulatory proteins called transcription factors (TFs) and *cis*-regulatory elements found on the promoters of regulated genes. Many external and internal signals, such as metabolite concentrations, oxygen, temperature, pH levels or surface contact, are integrated into regulatory networks through different sensing mechanisms in order to provide an appropriate transcriptional response allowing the bacterium to adapt effectively to the new environment. TFs are classified in several families depending on the structural similarities in their DNA-binding domains (DBDs) which can either stand alone or be present in multi-domain proteins containing variable arrangements with sensing or oligomerization domains (Ulrich *et al.*, 2005). This allows the integration of countless different input and output signals into regulatory networks. Although TFs are classified in defined families (Sanchez *et al.*, 2020), they each control specific sets of genes to coordinate bacterial responses, and the targets and functions of each newly identified TF cannot be inferred but need to be experimentally established. One-component systems (OCS) are the most diverse TFs that can directly sense intracellular or imported extracellular signals through their signal-sensing domain and transmit the signal, probably through a conformational change, to their DBD (Ulrich *et al.*, 2005). Here we explored one family of OCS in *Pseudomonas aeruginosa*, a gram-negative opportunistic human pathogen.

*P. aeruginosa* possesses a considerable metabolic versatility and one of the most complex regulatory networks found in the bacterial kingdom, with roughly 10% of all genes dedicated to transcription regulation (Stover *et al.*, 2000, Rodrigue *et al.*, 2000, Galan-Vasquez *et al.*, 2011). This regulatory network contains about 500 predicted TFs. While some regulatory pathways have been thoroughly studied, the vast majority of *P. aeruginosa* TFs are still uncharacterized. With the development of NGS-based methods dedicated to the genome-wide characterization of TFs, the view of complex interplays between and inside different families of TFs is starting to emerge (Huang *et al.*, 2019, Rajeev *et al.*, 2020). We recently characterized a transcriptional inhibitor, ErfA, recruited to down-regulate the expression of an horizontally transferred operon encoding a major virulence factor, ExlA, specifically in one *P. aeruginosa* lineage (Trouillon *et al.*, 2020). ErfA belongs to the large family of regulators with a XRE-like DBD that usually binds their DNA targets as homodimers. Besides its N-terminal DBD, ErfA possesses a C-terminal putative sensor domain with a “cupin” fold (Trouillon *et al.*, 2020). While ErfA was rewired to also regulate the horizontally acquired *exlBA* operon in *P. aeruginosa*, its main conserved target across *Pseudomonas* species is a metabolic operon, located adjacent to the *erfA* gene, with no role in bacterial virulence. ErfA was proposed to act as a sensory switch on this transcription unit in response to yet unknown conditions. While signal-sensing TFs are widely distributed in bacteria (Ulrich *et al.*, 2005), TFs sharing ErfA specific domain architecture have not been studied, raising the question of whether ErfA-like regulators could be common and important in metabolic and/or virulence regulation.

In this work, we focused on the subfamily of eight *P. aeruginosa* TFs that share ErfA architecture in order to examine their genome-wide regulatory targets. In addition to ErfA (Trouillon *et al.*, 2020), the family comprises two members with incompletely-defined regulons and five uncharacterized TFs. The combination of RNA-seq and DAP-seq approaches allowed us to define the regulons of six of these TFs. Each family member has specific targets ranging from one to twelve binding sites in promoters of genes related to small molecule uptake or processing. Regulators with XRE-cupin domains were found as local, specialized inhibitors that are widespread across the *Pseudomonas* genus. While many uncharacterized XRE-cupin TFs were identified, some species (such as *P. putida*) harbored up to ten of these regulators whereas others (i.e. *P. stutzeri*) possess only on, potentially reflecting different metabolic versatilities.

## RESULTS

### Eight TFs of the XRE-like family share similar architectures

The *P. aeruginosa* PAO1 genome encodes 19 proteins with a helix-turn-helix DNA-binding motif similar to that of the CI and Cro repressors of the phage λ that features the members of the XRE-like family (Wood *et al.*, 1990, Ohlendorf *et al.*, 1983) (Figure 1A). The XRE-like domains of about 60 residues adopt a well-characterized helical conformation with helices 2 and 3 involved in DNA binding (Figure 1B). The DBD was found alone or associated to either a peptidase S24 domain (AlpR and PrtR) or to a predicted sensor cupin domain at the C-terminus of proteins (Figure 1C). The versatile cupin domain folds into a 6-stranded β-barrel and is associated to a wide range of function (Dunwell *et al.*, 2004). In addition to ErfA (PA0225), seven TFs share the XRE-cupin architecture with only two having partially attributed functions: PA0535, PA1359, PA1884, PA2312, PA4499 (PsdR), PA4987 and PA5301 (PauR). Using targeted approaches, PsdR and PauR were previously shown to regulate dipeptide metabolism (Kiely *et al.*, 2008, Asfahl *et al.*, 2015) and polyamines metabolism (Chou *et al.*, 2013), respectively. By regulating independent metabolic pathways, these two TFs play an essential role for bacterial growth in their corresponding environmental conditions, which predicts potentially important functions for the five so far unstudied TFs of the family.

**FIGURE 1.**
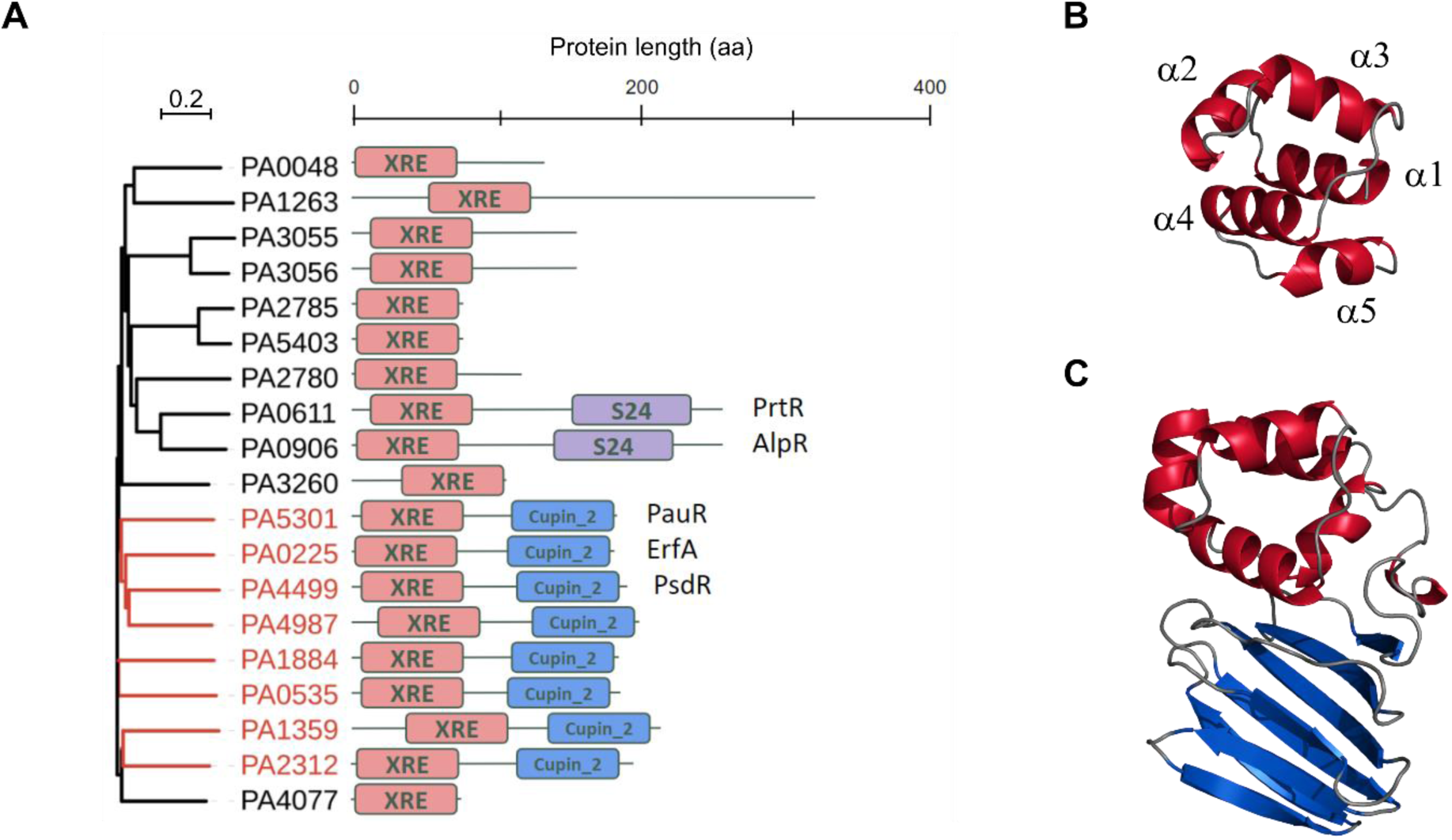
*P. aeruginosa* TFs belonging to the XRE family. (**A**) Maximum likelihood phylogenetic tree of the 19 proteins possessing a XRE DNA-binding domain in *P. aeruginosa* PAO1 (Winsor *et al.*, 2016). The leaves of the eight XRE-cupin regulators are colored in red in the tree. (**B**) Stereo ribbon representation of the model of the XRE DNA-binding domain of ErfA. The five alpha helices are annotated. (**C**) Stereo ribbon representation of the model of ErfA structure. The XRE domain is in red and the cupin domain in blue. Model prediction was done using the SWISS-Model tool (Biasini *et al.*, 2014) using the structure with PDB ID 1Y9Q as template.

### TFs of the XRE-cupin family are local, highly specialized repressors

To get a global view on target specificities and regulatory networks of *P. aeruginosa* XRE-cupin TFs, we undertook to determine the regulons of all members of this family, except for ErfA that we already comprehensively studied (Trouillon *et al.*, 2020). To that aim, the corresponding genes were deleted in the genome of PAO1 and the transcriptomes of the engineered mutants were compared to that of the wild-type strain by RNA-seq. In parallel, we expressed and purified the seven recombinant regulators, and their targets were identified *in vitro* on fragmented *P. aeruginosa* PAO1 genome by DAP-seq (Bartlett *et al.*, 2017, Trouillon *et al.*, 2020). Altogether, we determined transcriptomes and direct DNA targets for six of the seven regulators (Figure 2A, Tables S3 and S4). The PA2312 protein was less soluble and stable than any of the other proteins and was thus probably inactive and/or aggregated resulting in no significant peak detected in DAP-seq. In addition, the *PA2312* gene seems not expressed in the tested condition, as revealed by the few reads covering the CDS observed by RNA-seq in the wild-type strain. Overall, the family of XRE-cupin TFs regulates 39 genes in the PAO1 strain through direct binding to 19 genomic regions in total (Figure 2B).

**FIGURE 2.**
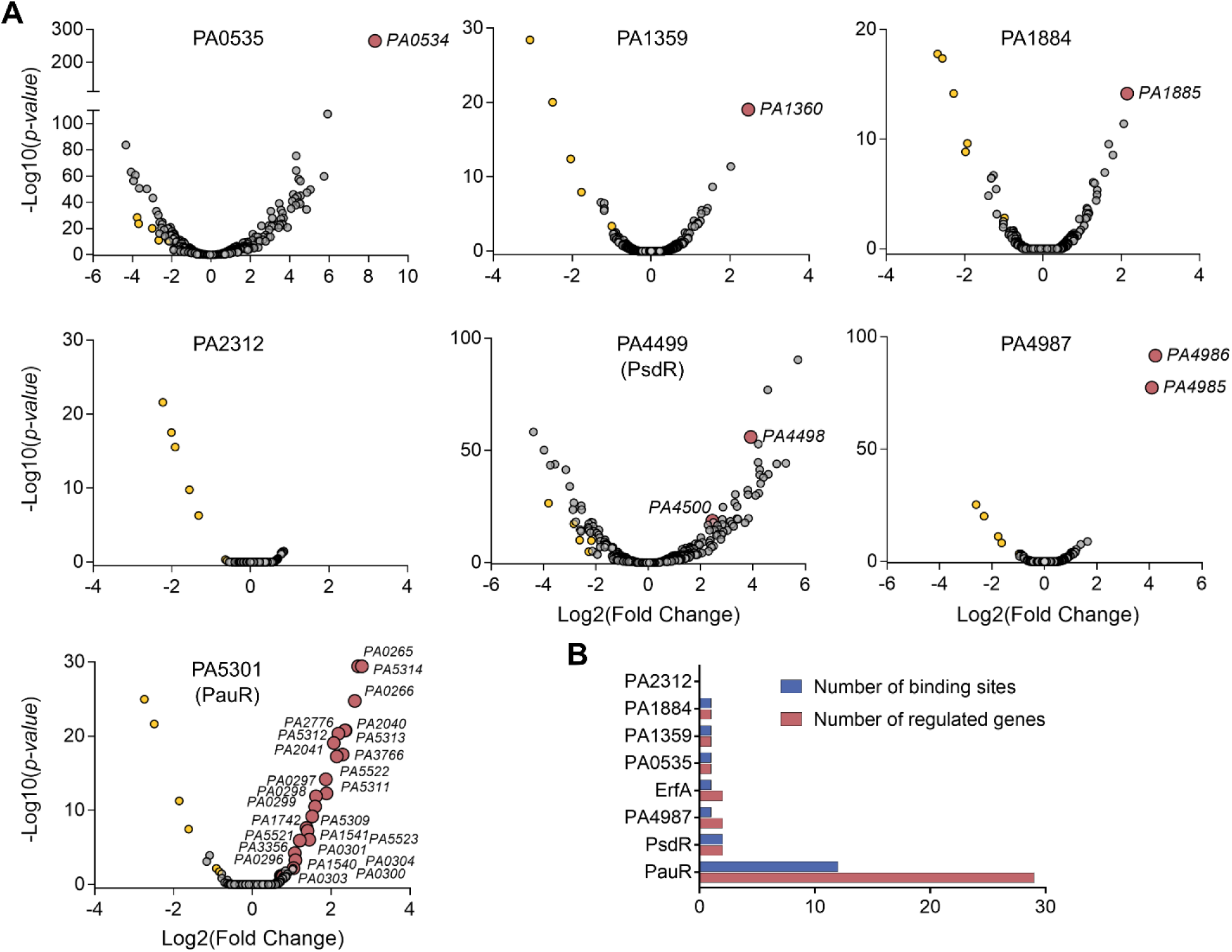
Determination of the regulons of the XRE-cupin regulators. (**A**) Volcano plots displaying the RNA-seq results of the genes differentially expressed in the respective XRE-cupin mutants versus parental strain PAO1. Genes for which a DAP-seq binding peak was identified in the promoter are represented by red circles and annotated with their gene ID. The five genes (yellow circles) found commonly downregulated in all mutants represent artefacts probably due to genetic manipulation. (**B**) Summary of number of regulatory targets per TF in *P. aeruginosa* PAO1.

In all cases, there was a strong correlation between *in vivo* and *in vitro* targets; the majority of DAP-seq peaks were centered on the promoters of genes that were also significantly dysregulated in RNA-seq. Most binding events occurred in the core promoter regions of regulated genes (Figure 3A) and each target gene was found upregulated in the mutant of the corresponding regulator (Figure 3B), showing that the XRE-cupin regulators are inhibitors of transcription. Our analyses confirmed the known regulon of PsdR and extended our knowledge on PauR, as discussed below. For most regulators, we found one or two direct targets, with the exception of PauR (PA5301) that directly regulates 29 genes (Figure 2B). Some genes were found slightly downregulated in all mutants with no corresponding binding sites found, probably due to experimental conditions or genetic manipulations (Figure 2A).

**FIGURE 3.**
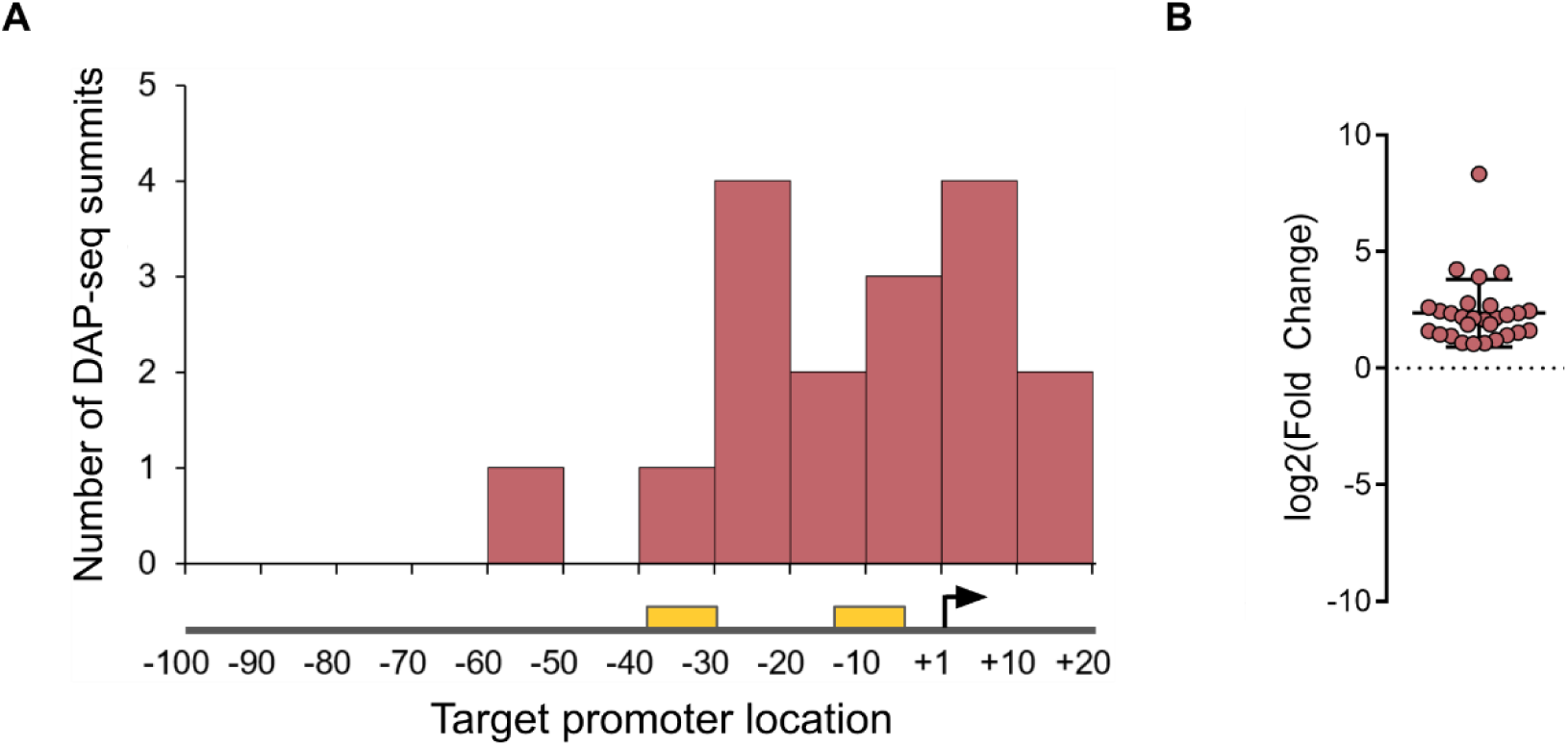
The XRE-cupin regulators are inhibitors of transcription. (**A**) Repartition of binding sites identified by DAP-seq within target promoters. RNA polymerase binding sites were either inferred from experimentally determined transcription start sites (Wurtzel *et al.*, 2012) or predicted using BPROM (http://softberry.com) if no data was available. The transcription start site is shown as a black arrow, and the −10 and −35 boxes as yellow rectangles. (**B**) RNA-seq expression fold changes of target genes in regulators mutants compared to the wild-type strain.

At least one regulatory target per TF was further verified by RT-qPCR and electrophoretic mobility shift assays (EMSA) and in all cases the RNA-seq and DAP-seq results were confirmed by these targeted approaches (Figure 4).

**FIGURE 4.**
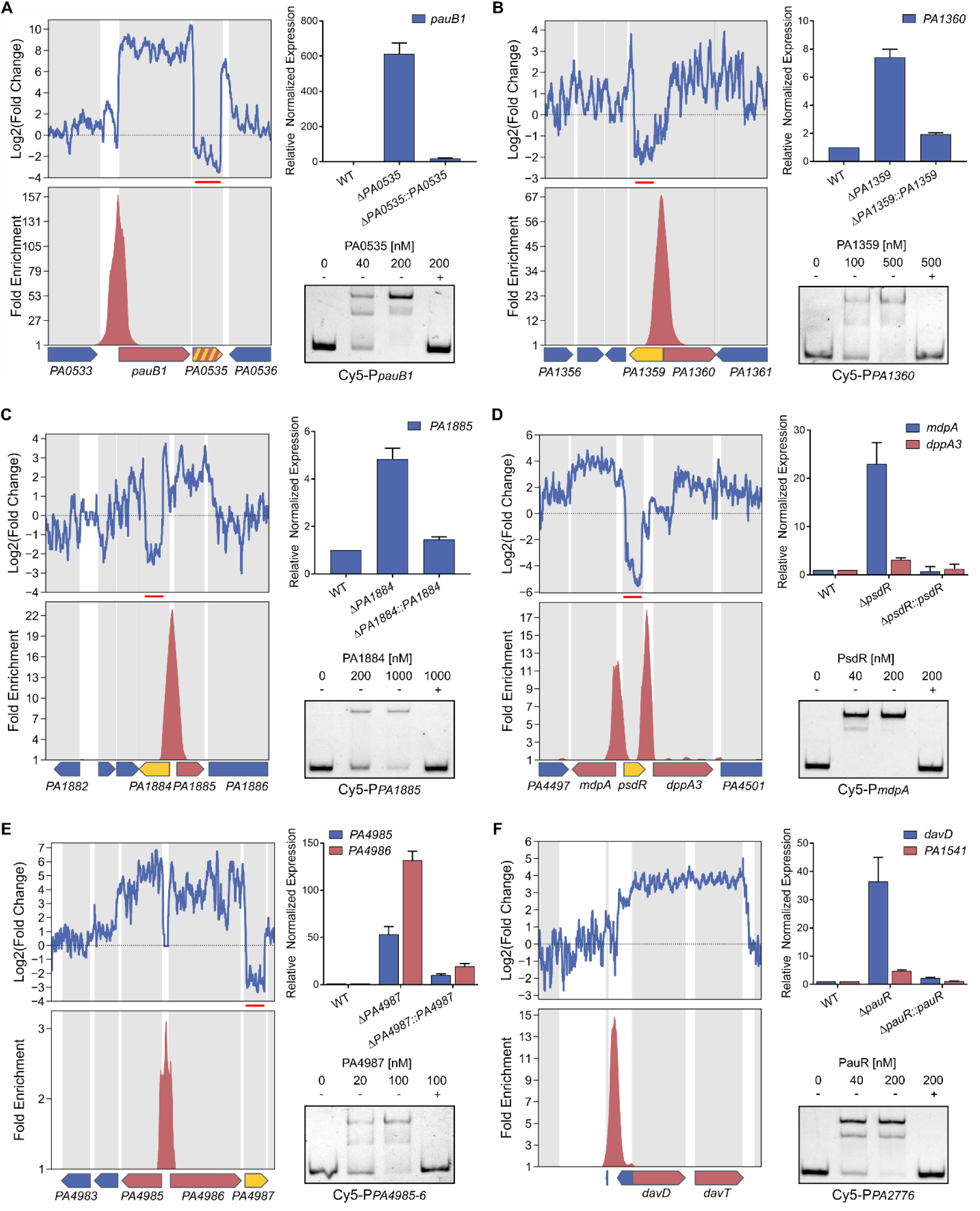
RT-qPCR and EMSA confirm the genome-wide results. Selected TFs and targets were PA0535 and *pauB1* (*PA0534*) (**A**), PA1359 and *PA1360* (**B**), PA1884 and *PA1885* (**C**), PsdR and *mdpA* (*PA4498*) and *dppA3* (*PA4450*) (**D**), PA4987 and *PA4985*-*PA4986* (**E**), PauR and *davD* (*PA0265*), *PA1541* (by RT-qPCR), and *PA2776* (by EMSA) (**F**). Upper left panels: local RNA-seq read abundance fold changes in the corresponding mutants compared to parental strain. Red lines show the deleted region in the mutant strain. Lower left panels: local fold enrichments in the corresponding regulator obtained by DAP-seq compared to negative controls. Target genes are shown as red arrows, genes encoding the studied TF as yellow arrows and others as blue arrows. Suspected autoregulation are denoted with dashed yellow-red arrows. Upper right panels: RT-qPCR showing the regulation of the target genes in the corresponding regulatory mutants and complemented strains. Experiments were performed in triplicates and normalized to the *rpoD* transcripts. Error bars indicate the SD. Lower right panels: EMSA on target binding sites. Recombinant XRE-cupin-His_10_ proteins were incubated with 0.5 nM Cy5-labeled probes for 15 min before electrophoresis. For competition assays, excess of unlabeled probes (100 nM) is denoted ‘+’.

Strikingly, all the XRE-cupin TFs bind to at least one intergenic region directly adjacent to their own gene (Figure 5). PauR strikes as an outlier with its 29 regulated genes scattered across twelve different genomic locations, while all other XRE-cupin repressors have one or two targets, always found in the direct vicinity of their own genes, showing that they are local, specialized regulators that form local functional units with their regulated genes. Horizontal gene transfer (HGT) is thought to shape the evolution of these gene groups and their regulatory relationships, and might be at the origin of their differentiation from one another (Hershberg *et al.*, 2005, Price *et al.*, 2008).

**FIGURE 5.**
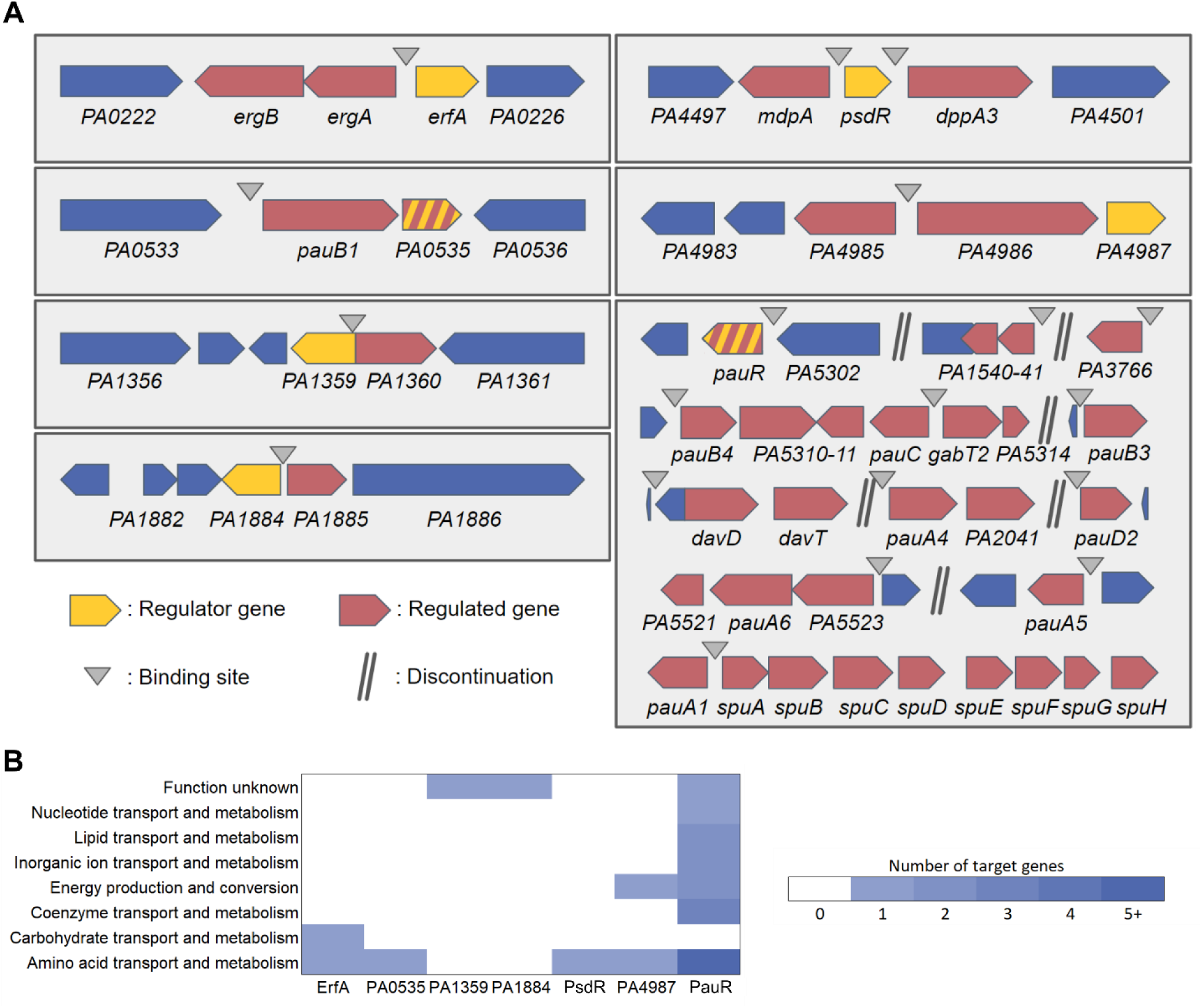
The complete regulons of the XRE-cupin regulators. (**A**) Schematic views of the local targeted regions of the seven XRE-cupin with determined regulons in *P. aeruginosa* PAO1. (**B**) Functional annotation of XRE-cupin regulatory targets. COG functional annotations were retrieved from PAO1 genome on the *Pseudomonas* database (Winsor *et al.*, 2016) for all target genes.

Based on DAP-seq peaks, DNA-binding motifs could be generated for PsdR and PauR, which have more than one binding site (Figure S1). These motifs exhibit a palindromic architecture, as the previously determined ErfA consensus (Trouillon *et al.*, 2020), supporting DNA binding of XRE-cupin TFs as dimers. Mutations introduced into the binding site prevented PauR binding, further validating the inferred consensus (Figure S1). Interestingly, mutation of only half the palindromic sequence recognized by PsdR led to a faster mobility fragment than that obtained with the wild-type sequence. This might correspond to the binding of one monomer to the mutated probe and supports the model in which the usual two-band shift observed with most XRE-cupin TFs corresponds to the binding of a monomer (lower, fainter band) and of a dimer (higher, stronger band) at higher concentration (Figures 4 and S1). The three determined binding consensus are quite different, probably due to differences in the amino acid composition of their DNA binding interface observed among the TF family (Figure S2). This supports their different DNA binding preferences and regulon specificities, as no common targets were revealed for any of the regulators.

Altogether, we defined the regulons of seven TFs, completing our knowledge of the XRE-cupin TF family in *P. aeruginosa* (Figure 5A). Our results show that all the members of the family are inhibitors of transcription, most likely acting by occluding the promoter and preventing RNA polymerase recruitment. Despite of their common features (domain architecture, local action, inhibitory function), each XRE-cupin TF targets specific regulons without overlapping roles.

### XRE-cupin TFs regulate specific metabolic pathways

To determine the functions of each TF, we scrutinized the roles of their regulons in details (Figures 5A and 5B). That of ErfA was already mentioned and encompasses the *ergAB* operon in PAO1, playing a putative role in amino acid or carbohydrate metabolism (Trouillon *et al.*, 2020). PA0535 has only one target: it inhibits the *pauB1* gene and potentially its own expression, the two genes being predicted as an operon (Winsor *et al.*, 2016) (Figures 4A and 5A). PauB1 is one of the four redundant PauB proteins, which are FAD-dependent oxidoreductases working in the complex γ-glutamylation pathway required for polyamine utilization in *P. aeruginosa* (Chou *et al.*, 2013, Luengo & Olivera, 2020). Although expression of its gene was shown to be increased in presence of putrescine (Chou *et al.*, 2008), PauB1 is required for cadaverine catabolism (Chou *et al.*, 2013). We thus identified PA0535 as the missing regulator of the γ-glutamylation pathway, most of the genes being under the control of PauR, or BauR (Luengo & Olivera, 2020).

Indeed the key regulator PauR was shown to bind *in vitro* to eight sites and thus to potentially affect expression of 18 genes (Chou *et al.*, 2013). As mentioned above, our results confirmed all known DNA targets of PauR and RNA-seq provided further information concerning the extent of gene expression perturbation resulting from the TF binding. Two previously reported PauR binding sites were predicted to control the expression of six genes (Chou *et al.*, 2013), while our genome-wide expression data show that they actually impact the expression of 13 genes, *pauA1* and the *spuABCDEFGH operon* for one, and the two divergently transcribed two-gene operons, *pauC*-*PA5311* and *gabT2*-*PA5314* for the other. Furthermore, we also extended the PauR regulon by identifying four additional binding sites, one located upstream of *pauR* gene, strongly supporting an autoregulation mechanism. Three new identified targets were the *PA3766* gene encoding a probable amino acid/polyamine transporter and two operons, *davD-davT* and *PA1541-40*, which we confirmed by RT-qPCR (Figure 4F). While *PA1541* encodes a probable transporter, the DavD and DavT proteins are enzymes involved in the conversion from 5-aminovalerate (AMV) to glutarate (Luengo & Olivera, 2020). Cadaverine is converted into AMV through the γ-glutamylation pathway, showing that PauR regulation extends beyond this metabolic pathway to further downstream steps. The PA1359 regulator negatively controls the expression of the *PA1360* gene (Figure 4B), coding for a putative drug/metabolite transporter similar to the threonine exporter RhtA in *E. coli* (Livshits *et al.*, 2003). PA1884 has also only one target, the *PA1885* gene (Figure 4C), which product is a putative acyltransferase with a GNAT (Gcn5-related N-acetyltransferases) domain, which could either confer antibiotic resistance or have potential metabolic functions.

Even if *PA1885* is predicted in operon with the downstream *polB* gene (Winsor *et al.*, 2016), RNA-seq data indicated no difference on this gene, excluding a control by the inhibitor and suggesting the existence of a *polB*-specific promoter (Figure 4C). The PsdR regulator was already identified as the repressor of *dppA3* and *mdpA* (Kiely *et al.*, 2008), involved in the uptake and utilization of small peptides, our results completes the scheme by demonstrating that the control is direct and involves two binding sites surrounding the *psdR* gene (Figure 4D). DppA3 is a substrate-binding protein delivering tripeptides/dipeptides to the ABC transporter DppBCDF (Pletzer *et al.*, 2014), and MdpA is a metallo-dipeptidase involved in their processing (Kiely *et al.*, 2008). Finally, PA4987 has one binding site in the intergenic sequence of *PA4985* and *PA4986*, downregulating expression of both genes as confirmed by RT-qPCR (Figure 4E). Although predicted as an operon with *PA4985*, we did not observe any effect on *PA4984* expression (Figure 4E). PA4986 is a putative oxidoreductase and PA4985 a possible periplasmic spermidine/putrescine-binding protein. Overall, the XRE-cupin regulators seem to regulate functions related to specific amino acids or small molecule uptake or processing (Figure 5B).

Based on our knowledge of their different regulons, we attempted to further investigate the XRE-cupin TFs roles through two global phenotypic assays. We first took advantage of the *Galleria mellonella* infection model frequently used to assess overall bacterial fitness and virulence (Tsai *et al.*, 2016, Andrejko *et al.*, 2014, Hernandez *et al.*, 2019). No significant difference could be observed in survival curves between larvae infected with wild-type and mutant strains (Figure S3). Therefore, the overexpression of genes regulated by the XRE-cupin family of repressors did not provide *P. aeruginosa* any advantage or disadvantage in this simple animal model. On the other hand, we assessed the strains antibiotics resistance to 24 different clinically relevant antibiotics and again found no differences (Figure S3). These results reflect the specificity of the XRE-cupin regulators and their functional units towards precise conditions. Indeed, *psdR* inactivation was shown to provide a growth fitness during proteolytic growth in caseinate medium, as this provides the dipeptide substrates for the derepressed dipeptides uptake and degradation (Asfahl *et al.*, 2015). Also, *mdpA* expression is induced in presence of X-pro dipeptides that might explain the faster growth in their presence due to their higher metabolism (Kiely *et al.*, 2008). The fact that they do not affect fitness in global phenotypic assays illustrates that TFs of this family, along with their target genes, are niche-specific functional units allowing the bacteria to detect and respond to defined conditions.

It is hypothesized that XRE-cupin TFs are able to detect small signal molecules and detach from their DNA binding sites upon sensing in order to induce expression of the target genes. PauR was indeed shown to be involved in the derepression of genes of the γ-glutamylation pathway in presence of putrescine and cadaverine (Chou *et al.*, 2013). In addition, the *E. coli* PauR homolog PuuR is known to dissociate from its binding site upon direct sensing of putrescine (Nemoto *et al.*, 2012). In the same line, *mdpA* expression was shown to be induced in presence of different dipeptides, probably sensed by PsdR (Kiely *et al.*, 2008). In all these cases, the signals were metabolites that are part or the precursors of the regulated metabolic pathways. The cupin domain in these regulators is predicted as a signal-sensing domain and thus could be responsible for the specific sensing of each regulon-related metabolites, as it is often the case (Fernandez-Lopez *et al.*, 2015). The crystal structure of a XRE-cupin regulator from *Vibrio cholera* has been solved and shows a binding pocket in the cupin domain with bound D-methionine (PDB ID: 1Y9Q). The structure modelization of all eight *P. aeruginosa* XRE-cupin regulators also led to the identification of specific binding pockets in each cupin domain (Figure S4). Similarly, to what was found for their DNA-binding interfaces (Figure S2), each regulator exhibits a different amino-acid composition inside of this putative signal-sensing pocket. These variabilities at the two major functional regions of the proteins explain the difference in DNA- and signal molecule-binding preferences and may result from evolutionary processes involving HGT that led to the multiplication and differentiation of these regulators.

### XRE-cupin regulators are diverse and differently conserved between *Pseudomonas* species

As XRE-cupin TFs seem to regulate small individual metabolic pathways that could reflect the different environments encountered by the bacteria and their ability to adapt, we assessed the conservation of this family across the *Pseudomonas* genus. We first examined the conservation of the eight regulators studied here and found different conservation between *Pseudomonas* species (Figure 6A). Interestingly, PA1359 and PauR were conserved in nearly all strains, suggesting a more central role or ancestral origin. The six other regulators are less conserved between strains and species. We can notice that although PA2312 seems not to be active in the PAO1 strain, its gene is conserved within *P. aeruginosa* strains and is present in other species, underlining its physiological importance. Here again, the conservation of the TFs and their regulatory targets probably depends on the environments encountered by each species which might require or not the associated metabolic functional units.

**FIGURE 6.**
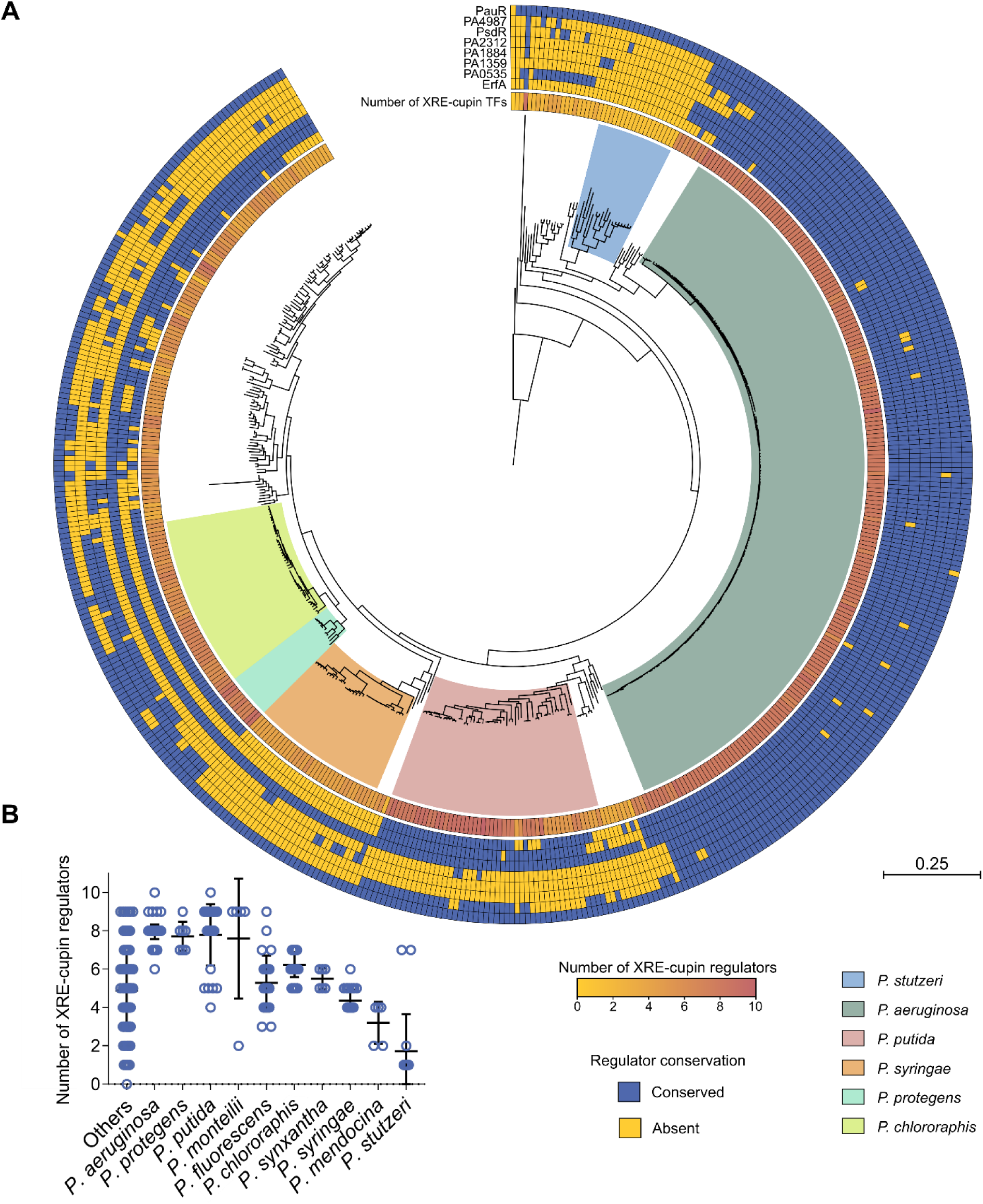
Phylogenetic analysis of the conservation of the XRE-cupin regulatory family across the *Pseudomonas* genus. (**A**) Maximum-Likelihood phylogenetic tree of 503 *Pseudomonas* complete genomes. The tree was generated from the multiple alignment of the concatenated sequences of 66 core genes for each strain with 100 bootstraps. The most represented species are delineated with a colored background. The number of XRE-cupin proteins detected through the Hidden Markov Model search is shown as the inner circle yellow-to-red heatmap. The outer circles show the results of homolog search by reciprocal best blast hit search for the eight regulators studied here. (**B**) Dot plot showing the distribution of number of XRE-cupin regulators per strain for each species. Species represented by less than five strains are grouped in the “Others” column.

To apprehend the importance of these regulators, we investigated the presence of other XRE-cupin TFs in all strains. Using all XRE-cupin sequences available on NCBI (*n* = 23,072) as templates, we created and validated a Hidden Markov Model for the automated detection of such regulators. The model was able to specifically detect all XRE-cupin TFs in PAO1 (Figure S5). Using this model, we screened nearly three million proteins retrieved from all *Pseudomonas* complete genomes for the presence of XRE-cupin regulators and identified 3,147 of them, with between zero and ten present per strain, depending on the strain or species (Figures 6A and 6B). Some species were found with only few XRE-cupin TFs, less than two or three on average for some of them as for example in *P. mendocina* or *P. stutzeri*. This could reflect the fact that these species are more niche-specific and may not encounter a wide variety of environments and thus need less of these optional metabolic response functional units. On the other hand, species like *P. aeruginosa, P. protegens* or *P. putida*, possess around eight and up to ten XRE-cupin regulators. All three of these species are known for their versatility and capacity to adapt to a wide array of environmental conditions. The total number of XRE-cupin regulators per strain often encompasses other TFs than the eight studied here, showing that much more regulators of this family are present across the *Pseudomonas* genus, probably associated to as many more different metabolic functional units. For instance, while only five of the eight TFs studied here are conserved in *P. putida*, most strains of this species possess eight or nine XRE-cupin TFs. This exemplifies the diversity of this family of TFs and shows that many species or strains have evolved new functional units including new XRE-cupin regulators to respond to the specific environmental niches they might encounter.

### XRE-cupin regulators control neighboring enzyme-coding and metabolism-related genes

Six out of the seven XRE-cupin TFs experimentally characterized here regulate at least one gene adjacent to their own gene, forming local functional units. To investigate whether the local target regulation is universal for XRE-cupin TFs across the *Pseudomonas* genus, we first investigated the conservation of their neighboring genes (Figure 7A). We found a high conservation of regulator/regulated genes pairs, highlighting the local nature of these functional units and the fact that they are exchanged by HGT as a group of genes between bacteria. To go further on the characterization of all XRE-cupin regulators, we assessed the functions of the neighbor genes of all 3,147 regulators identified here. Based on GO Terms annotations of all 6,294 direct genetic neighbors, the vast majority of them encodes either enzymes or small molecule transporters (Figure 7B). This result strongly corroborates the fact that local functional units comprising XRE-cupin regulators act as metabolic modules for the response to the presence of specific metabolites.

**FIGURE 7.**
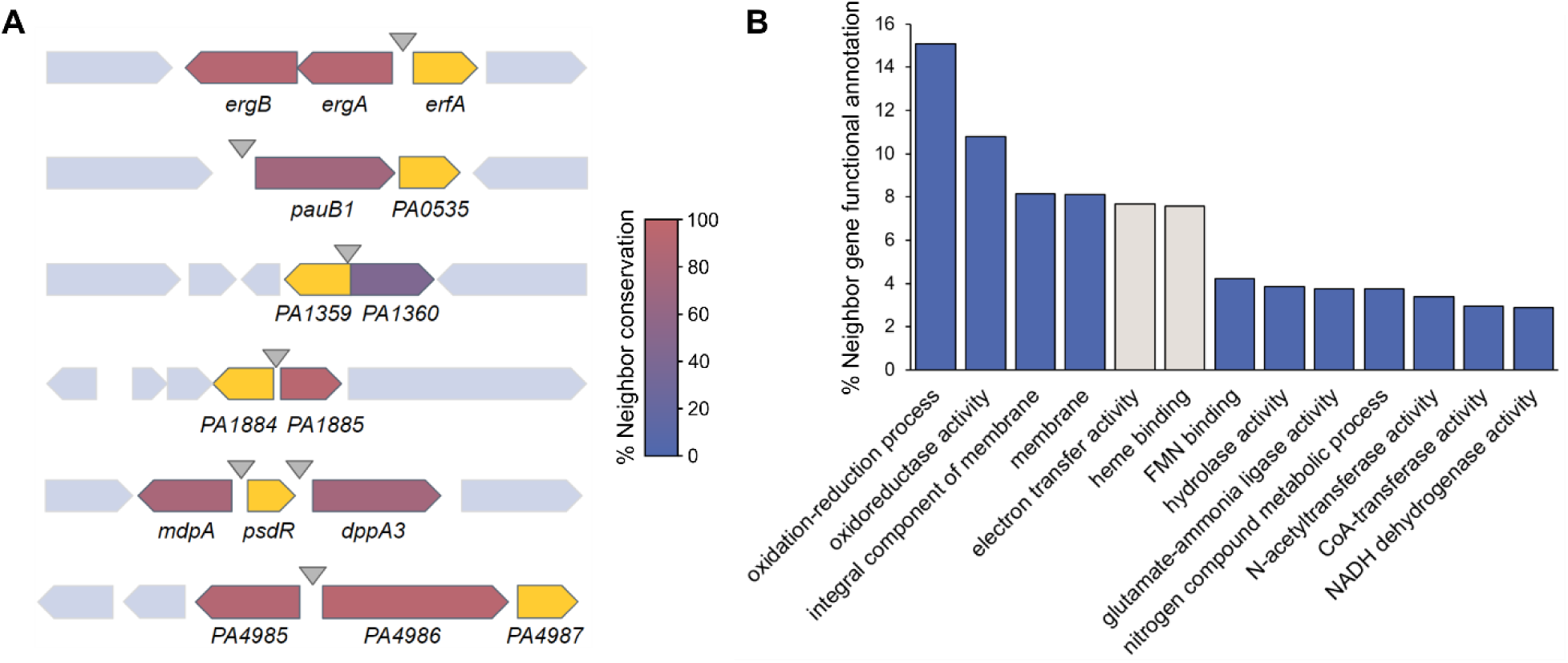
Gene synteny and genetic environment of the XRE-cupin regulators. (**A**) Conservation of XRE-cupin neighboring target genes. Regulator-coding genes are shown in yellow. Target genes are colored depending on how often they were found as conserved neighbors of their associated XRE-cupin TF. (**B**) Histogram showing the proportion of the most represented (>2%) GO functional annotations. Functional annotations were obtained from Pfam results of Interproscan search on 6,294 genes neighboring XRE-cupin regulators from 503 *Pseudomonas* genomes. Two categories are shown in grey as they correspond to the *cycB* gene, a conserved neighbor of *pauR* not regulated by PauR, and thus are not representing XRE-cupin regulatory targets.

## DISCUSSION

In the present study, we characterized a family of eight TFs sharing the same domain architecture (XRE-cupin TFs) in *P. aeruginosa* and identified their regulatory targets. The XRE-cupin family members are exclusive inhibitors of the transcription of metabolism-related genes probably needed in defined, specific conditions. The current model is that these TFs act as regulatory switches that keep the transcription of their neighboring target genes down until their products are needed for a given metabolic pathway. Upon sensing of the precursor metabolite of the regulated pathway through their cupin domain, they disengage from the DNA, in order to allow transcription of the genes coding for enzymes or transporters that will process the sensed molecule. This could include catabolism for nutrient source, as seen for PsdR, or detoxification via export of sensed molecules, as predicted for PA1359.

In most cases, the target genes were found located in the direct vicinity of the regulator gene, revealing the local nature of these regulatory interactions. Neighbor regulation is a common feature in bacteria, which has been explained by different models (Hershberg *et al.*, 2005, Lawrence, 2003). Genes that are functionally related, including regulators and regulated targets, tend to regroup together on the chromosome in order to be a complete functional unit when transferred by HGT. Different types of neighbor regulation are known (Hershberg *et al.*, 2005), three of which were found in this study; simple neighbor regulation and co-regulation of neighbors in both *cis* and *trans*, with some instances of autoregulation. In all cases, the total number of targets was small, allowing the formation of these local functional units. The very specific functions of the XRE-cupin regulons explain the small number of regulated genes and the fact that these units are local and probably easily transferrable gene clusters.

We found a high conservation of some of the studied TFs and their neighboring target genes across the *Pseudomonas* genus. These functional units may have been exchanged by HGT between bacteria that encounter similar environmental conditions. Additionally, a large number of different XRE-cupin regulators, each associated with their specific genetic neighborhood, were identified across the *Pseudomonas* genus. Out of the eight TFs studied here, none shared any common regulatory targets, and the two known sensed signals (dipeptides and polyamines) are different. This is explained by differences found at both DNA-binding and signal-sensing interfaces. Such differences are thought to happen through evolution after and during repurposing or duplication events and explain the presence of different XRE-cupin regulators in each species. The large number of this type of regulators illustrates their ancestral nature and their importance as condition-specific, transferrable means of metabolic diversification.

*Pseudomonas* species are famous for their wide metabolic versatility, encompassing many condition-specific metabolic pathways (Silby *et al.*, 2011). While the prediction of metabolism-related functions works well, experimental studies are still needed to pinpoint the exact pathways represented by uncharacterized genes. As illustrated by the target genes found here which mostly have only predicted functions, there is still a great need of phenotypic characterization of putative new metabolic pathways. Such studies would complete our knowledge on the mode of action of the XRE-cupin family of regulators and help understand both the function and sensed signals of these class of TFs. We also believe that family-wide and inter-species studies of TFs should become more common in order to be able to decipher global regulatory networks, especially in bacteria where they are still scarce.

## MATERIALS AND METHODS

### Bacterial strains

The bacterial strains used in this study are listed in Supplementary Table 1. *P. aeruginosa* and *E. coli* strains were grown in Lysogeny Broth (LB) at 37°C under agitation (300 rpm). *P. aeruginosa* strains were selected on Pseudomonas Isolation Agar (PIA). Antibiotics for *P. aeruginosa* were added when needed at the following concentrations: 200 μg/ml carbenicillin, 200 μg/ml gentamicin and 200 μg/ml tetracycline, and for *E. coli*: 100 µg/ml ampicillin, 50 μg/ml gentamycin and 10 μg/ml tetracycline.

### Plasmids and genetic manipulations

Plasmids and primers are listed in the Supplementary Tables S1 and S2, respectively. For overproduction of recombinant His10-tagged proteins, each gene sequence was amplified by PCR using PAO1 genomic DNA as a matrix and appropriate primer pairs, then integrated by SLIC in pET-52b cut with *Nco*I-*Sac*I and sequenced.

To generate *P. aeruginosa* deletion mutants, upstream and downstream flanking regions of each gene were amplified using appropriate primer pairs (sF1/sR1 and sF2/sR2 for each construct). The two resulting, overlapping fragments were then cloned into *Sma*I-cut pEXG2 by Sequence- and Ligation-Independent Cloning (SLIC) (Li & Elledge, 2007) and sequenced.

For complementation of *PA1359, PA1884, PA4499* and *PA5301* mutants, a fragment encompassing the promoter (around 500 bp upstream from the start codon) and the CDS was amplified by PCR and cloned by SLIC into the mini-CTX1 plasmid cut by *Sma*I. When the gene was predicted as the second gene of an operon (like *PA0535, PA2312* and *PA4987*) (Winsor *et al.*, 2016), the operon promoter was directly fused to the regulators gene. To do so, two fragments were generated, the upstream fragment carrying around 500 bp upstream and the 3 first codons of the first gene of the operon, and the second one the few last codons and the entire CDS of the gene of interest. Then the two overlapping fragments were cloned by SLIC in pEXG2 and sequenced.

All mini-CTX1- and pEXG2-derived plasmids were transferred by triparental mating into *P. aeruginosa*, using the helper plasmid pRK600. For allelic exchange, merodiploids resulting from cointegration events were selected on PIA plates containing gentamicin. Colonies were then plated on NaCl-free LB agar plates containing 10% sucrose to select for the loss of plasmid. The resulting sucrose-resistant strains were checked for gentamicin sensitivity and the mutant genotypes were determined by PCR. For complementation using mini-CTX1-derived plasmids, bacteria with *att* site-inserted plasmid were selected on PIA plates containing tetracycline, then they were cured from the mini-CTX1 backbone by excising FRT cassette with pFLP2 plasmid as previously described (Hoang *et al.*, 1998).

### Structure modelling

Protein sequences from the eight *P. aeruginosa* XRE-cupin TFs were aligned against the RCSB Protein Data Bank (Berman et al., 2000) in order to find an experimentally obtained 3D structure to use as a template for modeling. The 1.90Å 3D structure of a XRE-cupin transcription factor from *Vibrio cholerae* (PDB id: 1Y9Q) was used as template as it permitted the best modelling confidence using the SWISS-MODEL tool (Biasini *et al.*, 2014).

### Protein purification

All the pET-52b plasmids were transformed into *E. coli* BL21 Star (DE3). For protein overproduction, overnight cultures were diluted to an OD_600_ of 0.05 in LB medium containing 100 µg/ml ampicillin and expression induced at an OD_600_ of 0.6 with 1 mM IPTG. After 3h of growth at 37°C, bacteria were harvested by centrifugation at 6,000 g for 10 min at 4°C and resuspended in L Buffer (25 mM Tris-HCl, 500 mM NaCl, 10 mM Imidazole, 1 mM PMSF, 5 % Glycerol, pH 8, containing Roche protease inhibitor cocktail). Bacteria were then lysed by sonication. After centrifugation at 66,500 g for 30 min at 4°C, the soluble fraction was directly loaded onto a 1-ml nickel column (Protino Ni-nitrotriacetic acid [NTA]; Macherey-Nagel). The column was washed with Wash Buffer (50 mM Tris-HCl, 500 mM NaCl, 5 % Glycerol, pH 8) containing increasing Imidazole concentrations (20, 40, and 60 mM), and proteins were eluted with 200 mM Imidazole. Aliquots from the peak protein fractions were analyzed by SDS-PAGE, and the fractions containing the proteins of interest were pooled and dialyzed against ErfA Buffer (50 mM Tris-HCl, 250 mM NaCl, 50 mM KCl, 10 % Glycerol, 0.5 % Tween20, pH 7).

### DAP-seq experimental procedure, sequencing and data analysis

DAP-seq was carried out in triplicates on PAO1 genomic DNA exactly as previously described (Trouillon *et al.*, 2020). Sequencing was performed by the high-throughput sequencing core facility of I2BC (http://www.i2bc.paris-saclay.fr) using an Illumina NextSeq500 instrument. Approximately 8 million single-end reads per sample were generated on average with >90% of reads uniquely aligning to the PAO1 genome. Data analysis was performed as previously described (Trouillon *et al.*, 2020).

### RNA isolation

*P. aeruginosa* strains were grown from overnight cultures diluted to an OD_600_ of 0.1 in 3 ml of fresh LB medium at 37°C under agitation in duplicate. Total RNA was isolated at an OD_600_ of 1.0 using hot phenol-chloroform extraction as previously described (Trouillon *et al.*, 2020).

### RNA-seq libraries construction, sequencing and data analysis

After RNA isolation and DNAse treatment, RNA sample quality was assessed on an Agilent Bioanalyzer, yielding RINs of 9 or higher. Then ribosomal RNAs were depleted using the RiboMinus Transcriptome Isolation Kit (Thermofisher) following manufacturer instructions. The cDNA libraries were then constructed from 50 ng of depleted RNA using the NEBNext Ultra II Directional RNA library prep kit following manufacturer instructions (NEB). Libraries were size-selected to 200-700 bp using SPRIselect beads and quality was assessed on the Agilent Bioanalyzer using High Sensitivity DNA chips. Sequencing were done on an Illumina NextSeq500 and approximately 12 million single-end reads per sample were generated. Data analysis was performed as previously described (Trouillon *et al.*, 2020).

### RT-qPCR

After total RNA isolation and DNase treatment, cDNA synthesis was carried out using 2 µg of RNA as previously described (Trouillon *et al.*, 2020). The experiments were performed with 3 biological replicates for each strain, and the relative expression of mRNAs was analyzed with the CFX Manager software (Bio-Rad) using the Pfaffl method relative to *uvrD* reference Cq values. Statistical analyses were performed by T-test. The sequences of primers are listed in Supplementary Table 2.

### Electrophoretic Mobility Shift Assay

Genomic DNA was used as a matrix to generate wild-type sequence probes. To amplify probes bearing mutagenized binding sites, three different matrices were created: the one with mutated binding site in P_*PA0534*_ resulted from the fusion of two overlapping fragments generated by primer pairs pEXG2-mut-PA0534-BS-sF1/sR1 and pEXG2-mut-PA0534-BS-sF2/sR2. PCR fragments with modified binding sites in P_*PA2776*_ and P_*mdpA*_ were produced by using primer pairs CTX-PA2776-lacZ-sF/PA2776-mut-lacZ-sR and CTX-mdpA-lacZ-sF/mdAmut-lacZ sR, respectively. Then all fluorescent DNA probes were generated by PCR in two steps: a first PCR using specific primer pairs amplified the target DNA regions (83-96 bp) flanked by a 21 bp-region which is targeted in a second PCR by a single Cy5-labelled primer. The resulting labelled probes were purified on DNA Clean up columns (NEB) then incubated at 0.5 nM for 5 min at 37°C in EMSA Buffer (10 mM Tris-HCl, 50 mM KCl, 10 mM MgCl_2_, 10 % Glycerol, 0.1 mg/ml BSA, pH 8) containing 25 ng/µl poly(dI-dC). For competition assays, 100 nM unlabelled DNA probes (200 fold excess) were incubated with the labelled probes. Recombinant proteins were added at the indicated concentration in a final reaction volume of 20 µl and incubated for an additional 15 min at 25°C. Samples were then loaded on a native 8 % Tris-Acetate-EDTA (TAE) polyacrylamide gel and run at 100 V and 4°C in cold 1X TAE Buffer. Fluorescence imaging was performed using a Chemidoc MP.

### Phylogenetic analysis and XRE-cupin identification

All *Pseudomonas* complete genomes (*n*=503) were retrieved from the *Pseudomonas* Genome Database (Winsor *et al.*, 2016). The sequences from 66 core genes were concatenated for each genome and a multiple alignment was performed with MAFFT Galaxy version 7.221.3 (Katoh & Standley, 2013) using default settings. The resulting alignment was used to build a Maximum-Likelihood phylogenetic tree using MEGA X with 100 bootstraps which was visualized and annotated using iTOL v5 (Kumar *et al.*, 2018, Letunic & Bork, 2019). XRE-cupin homolog identification was performed by Reciprocal Best Blast Hit (RBBH) analysis (Cock *et al.*, 2015) on the European Galaxy server (Jalili *et al.*, 2020) using protein sequences from *P. aeruginosa* PAO1 against all protein sequences from the other 502 genomes.

For XRE-cupin identification, ErfA protein sequence was used to retrieve all protein sequences (*n*=23,072) with a XRE-cupin domain architecture (HTH_3 and Cupin_2) from the NCBI Conserved Domain Architecture Retrieval Tool (Marchler-Bauer *et al.*, 2015). A multiple alignment was performed on these sequences using MAFFT Galaxy version 7.221.3 (Katoh & Standley, 2013) with default settings. The resulting alignment was used to build a hidden Markov model using hmmbuild from HMMER (Potter *et al.*, 2018). All 2,862,589 protein sequences from the 503 genomes were then scanned through the model using hmmscan from HMMER (Potter *et al.*, 2018). A bimodal distribution of HMMER scores could be observed which led to the determination of a true positive XRE-cupin threshold (HMMER score >100), that was validated by functional prediction from both sides (Figure S5). All proteins above this threshold (*n*=3,147) were considered XRE-cupin regulators.

### Genetic neighbor functional prediction

Gene IDs from all proteins matching the HMM model were used to identify the direct upstream and downstream neighbor genes from all 503 genomes annotation files using Python 3.7. The corresponding 6,294 protein sequences were then used for functional prediction analysis using the Interproscan functional prediction tool (Jones *et al.*, 2014). Go annotations associated with Pfam predicted functions were then used for functional prediction.

## ACKNOWLEDGMENTS

We are grateful to Peter Panchev, Yvan Caspar, Eric Faudry and Viviana Job for their help with *G. mellonella* experiments, antibiotics resistance assays, biochemistry and structural modelling advices, respectively. This work was supported by grants from Agence Nationale de la Recherche [ANR-15-CE11-0018-01], the Laboratory of Excellence GRAL financed within the Grenoble Alpes University graduate school (Ecoles Universitaires de Recherche) CBH-EUR-GS [ANR-17-EURE-0003], and the Fondation pour la Recherche Medicale [Team FRM 2017, DEQ20170336705]. Julian Trouillon received a Ph.D. fellowship from French Ministry of Education and Research. We further acknowledge support from CNRS, INSERM, CEA, and Grenoble Alpes University.

## AUTHOR CONTRIBUTIONS

Conceived and designed the experiments: J.T., I.A., S.E. Performed the experiments: J.T., M.R., V.S., S.E. Analyzed the data: J.T., I.A., S.E. Wrote the initial draft: J.T., S.E. Manuscript finalization: J.T., I.A., S.E.

