## Supplementary Information for "Transcription inhibitors with XRE DNA-binding and cupin signal-sensing domains drive metabolic diversification in *Pseudomonas*"

### Supplementary methods

***Galleria mellonella* infection assay**. *G. mellonella* larvae infections were performed as previously described (Trouillon *et al.*, 2020). Animals injected with sterile PBS served as a control for physical trauma. CFUs for each dilution were systematically counted from the insulin pen and were of 7±1 CFUs per injection. Twenty larvae were injected per condition. Infection development was followed for 24 h at 37°C and the animals were considered dead when not reacting to touch. Strains were independently randomized and blinded ensuring no bias in CFU spotting and counting, and in animal death counting which was also done by a different person. Statistical significance was assessed using a Log-rank test.

**Antibiotics resistance screening.** Antimicrobial susceptibility was assessed by broth microdilution using the automated BD Phoenix system (Becton Dickinson, France) according to the manufacturers' recommendations. In brief, colonies were suspended in BD Phoenix$^{TM}$ ID broth to obtain approximately a 0.5 McFarland ($1.5.10^8$ UFC/ml) suspension. Broth was then placed on BD Phoenix$^{TM}$ AP system that automatically adjusts the optical density and further dilutes the bacterial suspension in BD Phoenix$^{TM}$ AST broth (a cation-adjusted formulation of Mueller-Hinton broth containing 0.01% Tween 80) to obtain a final bacterial concentration in the AST broth of approximately $5.10^5$ CFU/ml. A redox indicator was then added to the AST broth. Inoculated AST broth was then poured in BD Phoenix$^{TM}$ NMIC-417 panel, loaded into the BD Phoenix$^{TM}$ M50 instrument and incubated at 35˚C ± 1˚C. Results were interpreted by the BD EpiCenter™ system according to CASFM-EUCAST (European Committee on Antimicrobial Susceptibility Testing) 2019 V1 breakpoints.

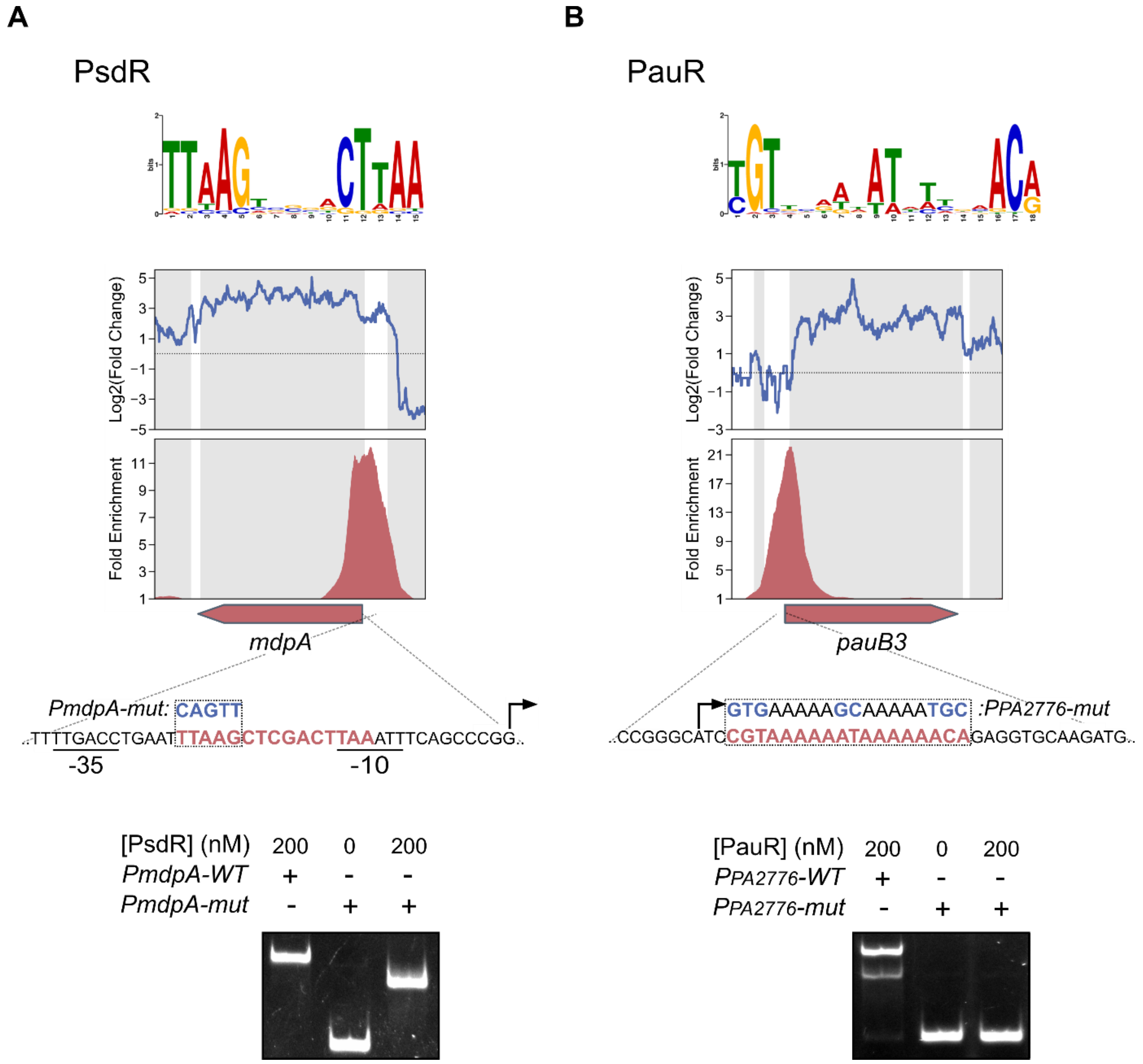


**FIGURE S1 : Validation of the DNA binding sites of PsdR and PauR.** For PsdR (**A**) and PauR (**B**), from upper to lower panels: (i) Enriched DNA motif obtained with MEME-ChIP in top DAP-seq peaks, (ii) local RNA-seq read abundance fold changes in the corresponding regulator mutants compared to the parental strain, (iii) local DAP-seq fold enrichments in the corresponding regulator experiments compared to negative controls, (iv) zoom view of the bound promoter region; mutated probes used in EMSA are shown with exchanged nucleotides in blue, and (v) EMSA of the corresponding region with either wild-type sequence or mutated probes. The mutation of half of the palindromic motif recognized by PsdR led to the shift of a lower band than with the wild-type probe, potentially representing the binding of only one monomer to the non-mutated half site. On the other hand, the mutation of the entire site for PauR led to a complete loss of binding.

**A**

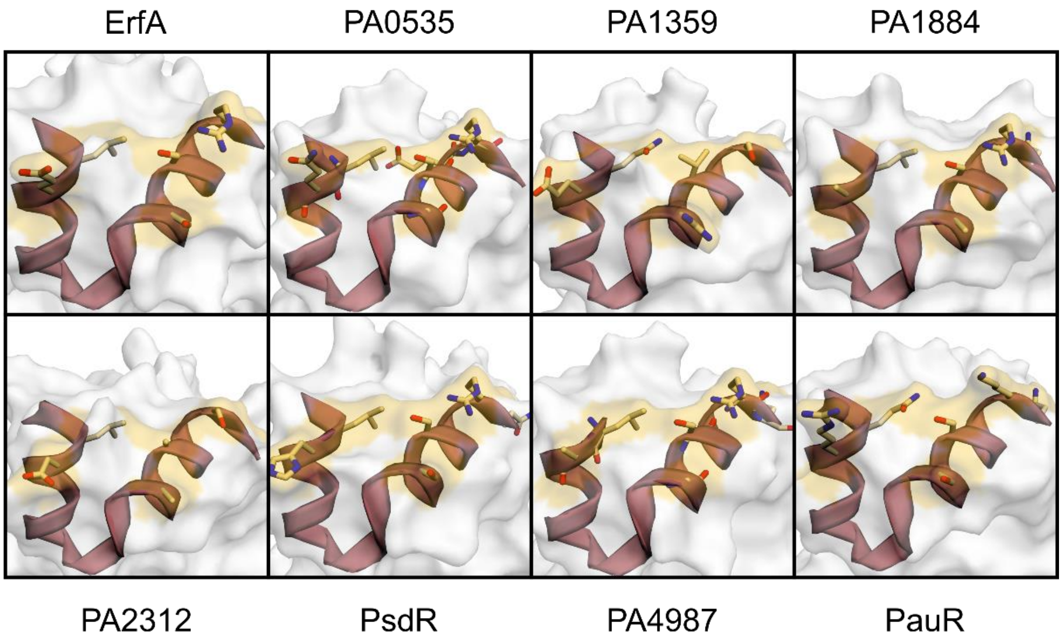


**B**

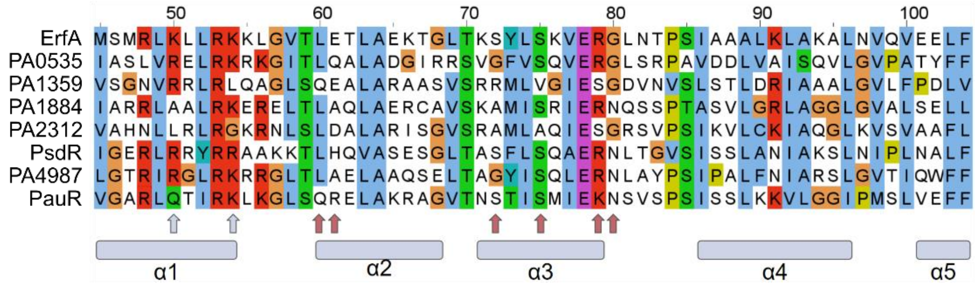


**FIGURE S2 : Predicted structures of DNA-binding interfaces of the eight XRE-cupin regulators.** (**A**) Stereo ribbon representation of the predicted DNA-binding interfaces of the eight XRE-cupin regulators. Model prediction was done using the SWISS-Model tool (Waterhouse *et al.*, 2018) and the structure with PDB ID 1Y9Q as template. Amino acids predicted to be involved in specific DNA-binding are depicted in yellow. (**B**) Sequence alignment of the eight XRE domains. The five alpha helices are denoted by grey rectangles. Amino acids predicted to be involved in specific and non-specific DNA-binding are shown by red and grey arrows, respectively.

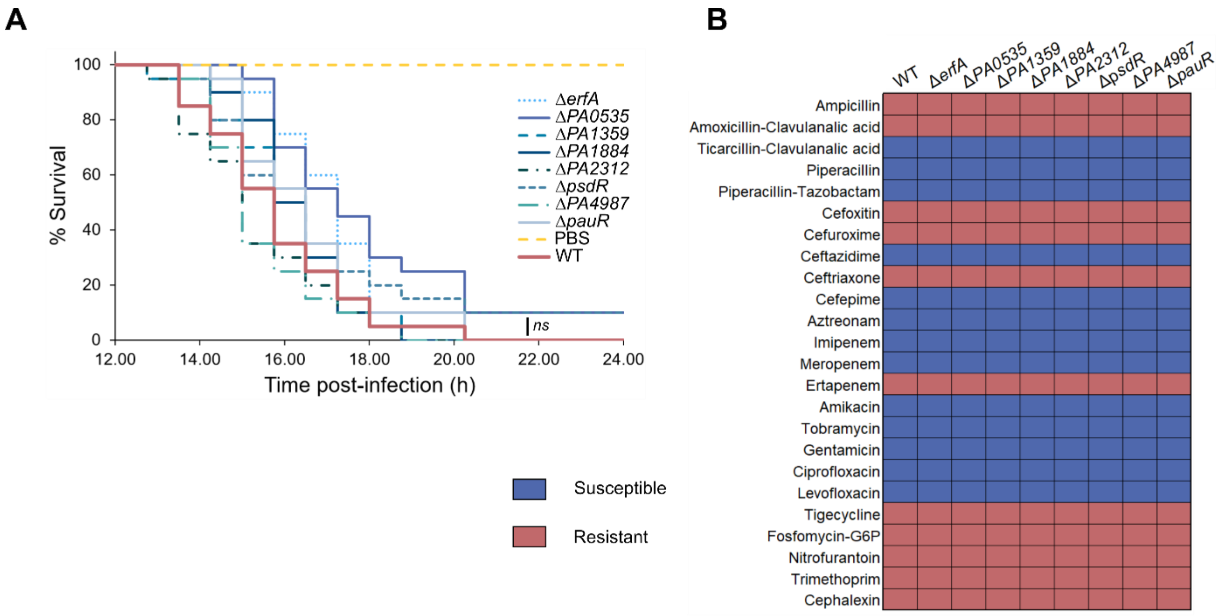


**FIGURE S3. The XRE-cupin regulons do not affect bacterial fitness in *G. mellonella*, nor antibiotics resistance.** (**A**) Survival curves of *Galleria mellonella* larvae infected with an average of 7-10 bacteria per larva. Twenty larvae were infected per strain. Significance testing was performed using log-rank. (**B**) Antibiotics resistance phenotypes of the XRE-cupin regulatory mutants. Resistance to 24 clinically relevant antibiotics was assessed in liquid using the automated BD Phoenix system.

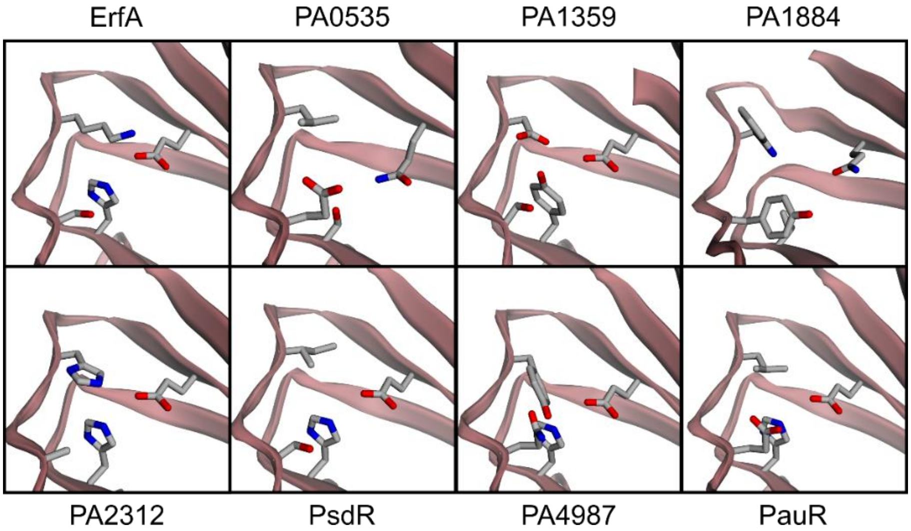


**FIGURE S4. Structure modeling of the XRE-cupin putative signal-sensing binding pocket.** Model prediction was done using the SWISS-Model tool (Waterhouse *et al.*, 2018) using the structure with PDB ID 1Y9Q as template. Inwards amino acids potentially involved in molecule binding are drawn.

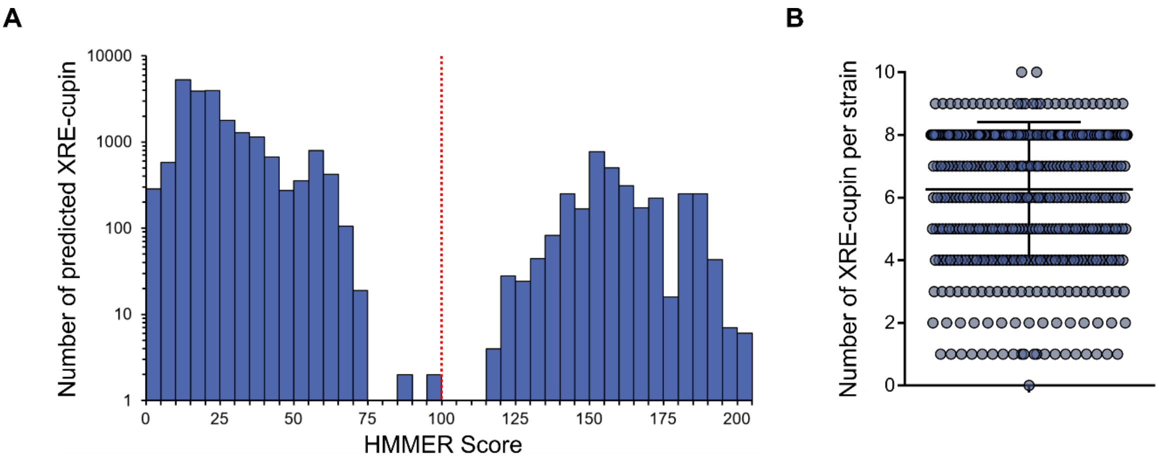


**FIGURE S5 : HMMER prediction of XRE-cupin regulators.** (**A**) Repartition of HMMER scores of all 23,974 proteins found to match the XRE-cupin HMM model. The bimodal distribution clearly delineates between false and true positives. All eight known *P. aeruginosa* XRE-cupin regulators had scores higher than 140 and all manually inspected predictions with scores < 100 were found not to be XRE-cupin proteins. Consequently, a threshold of > 100 (denoted by a red line) was used to consider true XRE-cupin regulators. (**B**) Repartition of the number of predicted XRE-cupin per strain.

**Table S1 | List of bacterial strains and plasmids used in this study**

| Strain or plasmid | Genotype or relevant properties | Reference/source |
|---|---|---|
| **Strains** | | |
| ***P. aeruginosa*** | | |
| PAO1 | Wound isolate, sequenced laboratory strain | J. Mougous |
| PAO1 Δ*erfA* | PAO1 with *erfA* (*PA0225*) deletion | (Trouillon *et al.*, 2020) |
| PAO1 Δ*erfA::erfA* | PAO1 Δ*erfA* with *erfA* in *attB* site | (Trouillon *et al.*, 2020) |
| PAO1 Δ*PA0535* | PAO1 with *PA0535* deletion | This work |
| PAO1 Δ*PA0535::PA0535* | PAO1 Δ*PA0535* with *PA0535* in *attB* site | This work |
| PAO1 Δ*PA1359* | PAO1 with *PA1359* deletion | This work |
| PAO1 Δ*PA1359::PA1359* | PAO1 Δ*PA1359* with *PA1359* in *attB* site | This work |
| PAO1 Δ*PA1884* | PAO1 with *PA1884* deletion | This work |
| PAO1 Δ*PA1884::PA1884* | PAO1 Δ*PA1884* with *PA1884* in *attB* site | This work |
| PAO1 Δ*PA2312* | PAO1 with *PA2312* deletion | This work |
| PAO1 Δ*PA2312::PA2312* | PAO1 Δ*PA2312* with *PA2312* in *attB* site | This work |
| PAO1 Δ*psdR* | PAO1 with *psdR* (*PA4499*) deletion | This work |
| PAO1 Δ*psdR::psdR* | PAO1 Δ*psdR* with *psrD* in *attB* site | This work |
| PAO1 Δ*PA4987* | PAO1 with *PA4987* deletion | This work |
| PAO1 Δ*PA4987::PA4987* | PAO1 Δ*PA4987* with *PA4987* in *attB* site | This work |
| PAO1 Δ*pauR* | PAO1 with *pauR* (*PA5301*) deletion | This work |
| PAO1 Δ*pauR::pauR* | PAO1 Δ*pauR* with *pauR* in *attB* site | This work |
| ***E. coli*** | | |
| TOP10 | Chemically competent cells | Invitrogen |
| BL21 Star (DE3) | $F^-$ *ompT hsdSB* ($rB^-$ $mB^-$) *gal dcm rne131*(DE3) | Invitrogen |
| **Plasmids** | | |
| pRK600 | Helper plasmid with conjugative properties ($Cm^R$) | (Kessler *et al.*, 1992) |
| pEXG2 | Allelic exchange vector ($Gm^R$) | (Rietsch *et al.*, 2005) |
| pEXG2-mut-PA0535 | pEXG2 carrying SLIC fragment for *PA0535* deletion ($Gm^R$) | This work |
| pEXG2-mut-PA1359 | pEXG2 carrying SLIC fragment for *PA1359* deletion ($Gm^R$) | This work |
| pEXG2-mut-PA1884 | pEXG2 carrying SLIC fragment for *PA1884* deletion ($Gm^R$) | This work |
| pEXG2-mut-PA2312 | pEXG2 carrying SLIC fragment for *PA2312* deletion ($Gm^R$) | This work |
| pEXG2-mut-PA4499 | pEXG2 carrying SLIC fragment for *PA4499* deletion ($Gm^R$) | This work |
| pEXG2-mut-PA4987 | pEXG2 carrying SLIC fragment for *PA4987* deletion ($Gm^R$) | This work |
| pEXG2-mut-PA5301 | pEXG2 carrying SLIC fragment for *PA5301* deletion ($Gm^R$) | This work |
| pFLP2 | Source of Flp recombinase ($Ap^R$) | (Hoang *et al.*, 1998) |
| mini-CTX1 | Site-specific integrative plasmid (*attP* site, $Tc^R$) | (Hoang *et al.*, 2000) |
| miniCTX-TrrnB-erfA | miniCTX1-TrrnB carrying *erfA* gene for complementation (*attP*, $Tc^R$) | (Trouillon *et al.*, 2020) |
| mini-CTX-PA0535 | mini-CTX1 carrying *PA0535* gene for complementation (*attP*, $Tc^R$) | This work |
| mini-CTX-PA1359 | mini-CTX1 carrying *PA1359* gene for complementation (*attP*, $Tc^R$) | This work |
| mini-CTX-PA1884 | mini-CTX carrying *PA1884* gene for complementation (*attP*, $Tc^R$) | This work |
| mini-CTX-PA2312 | mini-CTX1 carrying *PA2312* gene for complementation (*attP*, $Tc^R$) | This work |
| mini-CTX-PA4499 | mini-CTX1 carrying *PA4499* gene for complementation (*attP*, $Tc^R$) | This work |
| mini-CTX-PA4987 | mini-CTX1 carrying *PA4987* gene for complementation (*attP*, $Tc^R$) | This work |
| mini-CTX-PA5301 | mini-CTX1 carrying *PA5301* gene for complementation (*attP*, $Tc^R$) | This work |
| pET52b | Expression vector ($Ap^R$) | Novagen |
| pET52b-PA0535 | Expression vector of *PA0535-his10* ($Ap^R$) | This work |
| pET52b-PA1359 | Expression vector of *PA1359-his10* ($Ap^R$) | This work |
| pET52b-PA1884 | Expression vector of *PA1884-his10* ($Ap^R$) | This work |
| pET52b-PA2312 | Expression vector of *PA2312-his10* ($Ap^R$) | This work |
| pET52b-PA4499 | Expression vector of *PA4499-his10* ($Ap^R$) | This work |
| pET52b-PA4987 | Expression vector of *PA4987-his10* ($Ap^R$) | This work |
| pET52b-PA5301 | Expression vector of *PA5301-his10* ($Ap^R$) | This work |

**Table S2 | Primers used in this work**

| Name | Sequence (5' => 3') | Use |
|---|---|---|
| pEXG2-mut-PA0535-sF1 | **GGTCGACTCTAGAGGATCCCC**TCGTCCTCGACTACTTCCGC | *PA0535* deletion |
| pEXG2-mut-PA0535-sR1 | **GCGTATACCCAGAGCACCCTGTCA**GTCGTCGGCTACTTCGCTCA | *PA0535* deletion |
| pEXG2-mut-PA0535-sF2 | CAGGGTGCTCTGGGTATACG | *PA0535* deletion |
| pEXG2-mut-PA0535-sR2 | **ACCGAATTCGAGCTCGAGCCC**TCGGTGTCTATGTGGCGGTC | *PA0535* deletion |
| pEXG2-mut-PA1359-sF1 | **GGTCGACTCTAGAGGATCCCC** GGTGCGCACCGAGAGATAGA | *PA1359* deletion |
| pEXG2-mut-PA1359-sR1 | **CTGGCGAAGGCATGGAAGTCGTCA**GCCGGAGACATGTTCGAGGA | *PA1359* deletion |
| pEXG2-mut-PA1359-sF2 | CGACTTCCATGCCTTCGCCA | *PA1359* deletion |
| pEXG2-mut-PA1359-sR2 | **ACCGAATTCGAGCTCGAGCCC**ACGCTGGCTGAAGCCGTAGT | *PA1359* deletion |
| pEXG2-mut-PA1884-sF1 | **GGTCGACTCTAGAGGATCCCC**GTTGGTCGAGGAGCAGTTGC | *PA1884* deletion |
| pEXG2-mut-PA1884-sR1 | **CGTTGCAGGTGGAACGACTGCTCA**GATGCGCGAGATCATCGCCT | *PA1884* deletion |
| pEXG2-mut-PA1884-sF2 | GCAGTCGTTCCACCTGCAAC | *PA1884* deletion |
| pEXG2-mut-PA1884-sR2 | **ACCGAATTCGAGCTCGAGCCC**TGCCTGCTCATGCTGGGGTT | *PA1884* deletion |
| pEXG2-mut-PA2312-sF1 | **GGTCGACTCTAGAGGATCCCC**GCAGGTACGCGAGAGCCTTT | *PA2312* deletion |
| pEXG2-mut-PA2312-sR1 | **TAGAACAGGATCGAGTCGCCGTCA**GACGCGCTGGCCAATCAGAT | *PA2312* deletion |
| pEXG2-mut-PA2312-sF2 | CGGCGACTCGATCCTGTTCT | *PA2312* deletion |
| pEXG2-mut-PA2312-sR2 | **ACCGAATTCGAGCTCGAGCCC**CTGGCGATGATGGGCATGTC | *PA2312* deletion |
| pEXG2-mut-PA4499-sF1 | **GGTCGACTCTAGAGGATCCCC**TCACTGGGGTGCCATTGCAC | *PA4499* deletion |
| pEXG2-mut-PA4499-sF1 | **CAGGCCCATGGTGGTGATGGTTCA**TTCGCTGGCGACTTGGTGCA | *PA4499* deletion |
| pEXG2-mut-PA4499-sF1 | ACCATCACCACCATGGGCCT | *PA4499* deletion |
| pEXG2-mut-PA4499-sF1 | **ACCGAATTCGAGCTCGAGCCC**TAGTCGGTGCCGGTGGTGTA | *PA4499* deletion |
| pEXG2-mut-PA4987-sF1 | **GGTCGACTCTAGAGGATCCCC**GCTGTATGTGCGCGACCAGT | *PA4987* deletion |
| pEXG2-mut-PA4987-sF1 | **GGTGATCACCCAGAGCACCAGTCA**GGTGCCGAGGAAGTGCGTTT | *PA4987* deletion |
| pEXG2-mut-PA4987-sF1 | CTGGTGCTCTGGGTGATCAC | *PA4987* deletion |
| pEXG2-mut-PA4987-sF1 | **ACCGAATTCGAGCTCGAGCCC**TGCTGCTCGGCGATACCATG | *PA4987* deletion |
| pEXG2-mut-PA5301-sF1 | **GGTCGACTCTAGAGGATCCCC** TACAGCACCGAGCGTAGCCA | *PA5301* deletion |
| pEXG2-mut-PA5301-sF1 | **GTGGTGGCGCTGATCAAGCGTTCA**TTGCAGACGAGCACCGACGT | *PA5301* deletion |
| pEXG2-mut-PA5301-sF1 | ACGCTTGATCAGCGCCACCA | *PA5301* deletion |
| pEXG2-mut-PA5301-sF1 | **ACCGAATTCGAGCTCGAGCCC**CTTGCCGACGATGTCTTCGC | *PA5301* deletion |
| Comp-CTX-PA0535-sF1 | **GATATCGAATTCCTGCAGCCC**CAAGGAGATCGCCAAGGAACTG | *PA0535* complementation |
| Comp-CTX-PA0535-sR1 | CAGGAACATCGCGACCTCCGT | *PA0535* complementation |

| | | |
|---|---|---|
| Comp-CTX-PA0535-sF2 | **ACGGAGGTCGCGATGTTCCTG**CGTTTGCGCGACCTGTTCTGA | *PA0535* complementation |
| Comp-CTX-PA0535-sR2 | **TCTAGAACTAGTGGATCCCCC**CGACAGGGTAACGCGCCT | *PA0535* complementation |
| Comp-CTX-PA1359-sF | **GATATCGAATTCCTGCAGCCC**CGAAGAGGATGTACAGCGCCC | *PA1359* complementation |
| Comp-CTX-PA1359-sR | **TCTAGAACTAGTGGATCCCCC**TCCTCTGGAAGGCTGGCTCGA | *PA1359* complementation |
| Comp-CTX-PA1884-sF | **GATATCGAATTCCTGCAGCCC**GCTGCCGCTGACCGTGTCCA | *PA1884* complementation |
| Comp-CTX-PA1884-sR | **TCTAGAACTAGTGGATCCCCC**CCTCGACTGGTCGCCGGAG | *PA1884* complementation |
| Comp-CTX-PA2312-sF1 | **GATATCGAATTCCTGCAGCCC**TGAAGGCGATCTGCTGCGCC | *PA2312* complementation |
| Comp-CTX-PA2312-sR1 | GCGGTCCATGAGGCATTCCTC | *PA2312* complementation |
| Comp-CTX-PA2312-sF2 | **GAGGAATGCCTCATGGACCGC**CGGGCCTGACCGATGAACCT | *PA2312* complementation |
| Comp-CTX-PA2312-sR2 | **TCTAGAACTAGTGGATCCCCC**TCTTCGTCCATGGCAGCAAGG | *PA2312* complementation |
| Comp-CTX-PA4499-sF | **GATATCGAATTCCTGCAGCCC**TTCCTCCCAGGTGTGGATGGC | *PA4499* complementation |
| Comp-CTX-PA4499-sR | **TCTAGAACTAGTGGATCCCCC**GATGGGAAGGTGCGCTACGGA | *PA4499* complementation |
| Comp-CTX-PA4987-sF1 | **GATATCGAATTCCTGCAGCCC**CCTCGACCTGGACCAGGTCC | *PA4987* complementation |
| Comp-CTX-PA4987-sR1 | GAAATTCACGGTTGGCGTCCTT | *PA4987* complementation |
| Comp-CTX-PA4987-sF2 | **AGGACGCCAACCGTGAATTTC**AAGGCCGCCTGGAGCATCTGA | *PA4987* complementation |
| Comp-CTX-PA4987-sR2 | **TCTAGAACTAGTGGATCCCCC**CTGGCCCTGCATCGCGACAG | *PA4987* complementation |
| Comp-CTX-PA5301-sF | **GATATCGAATTCCTGCAGCCC**CGCTACAGCACCGAGCGTAG | *PA5301* complementation |
| Comp-CTX-PA5301-sR | **TCTAGAACTAGTGGATCCCCC**CTTGCGGCATTAGGTCGCAGC | *PA5301* complementation |
| pET52b-PA0535-sF | **TTAAGAAGGAGATATACC**ATGAGCGAAGTAGCCGACGA | *PA0535* overexpression |
| pET52b-PA0535-sR | **CTACCGCGTGGCACCAGAGCGAG**GGCGTATACCCAGAGCACC | *PA0535* overexpression |
| pET52b-PA1359-sF | **TTAAGAAGGAGATATACC**ATGGAAGAGGTCCGGCAGTG | *PA1359* overexpression |
| pET52b-PA1359-sR | **CTACCGCGTGGCACCAGAGCGAG**GCTCACCACGTTGCGGGTG | *PA1359* overexpression |

| | | |
|---|---|---|
| pET52b-PA1884-sF | **TTAAGAAGGAGATATACC**ATGGATATCGACGAACTGATTG | *PA1884* overexpression |
| pET52b-PA1884-sR | **CTACCGCGTGGCACCAGAGCGAG**GTCGTGGAGGATCACCAGC | *PA1884* overexpression |
| pET52b-PA2312-sF | **TTAAGAAGGAGATATACC**ATGCATACCGAACCCGATGATC | *PA2312* overexpression |
| pET52b-PA2312-sR | **CTACCGCGTGGCACCAGAGCGAG**GTCCAGGCGCTCCGGGTA | *PA2312* overexpression |
| pET52b-PA4499-sF | **TTAAGAAGGAGATATACC**ATGACCGTAGACCGCATCGG | *PA4499* overexpression |
| pET52b-PA4499-sR | **CTACCGCGTGGCACCAGAGCGAG**GGGCGTCGGATGGTCGTC | *PA4499* overexpression |
| pET52b-PA4987-sF | **TTAAGAAGGAGATATACC**ATGCCCCGCCCGTCACCG | *PA4987* overexpression |
| pET52b-PA4987-sR | **CTACCGCGTGGCACCAGAGCGAG**GAACGTCGGCGGGGTGATC | *PA4987* overexpression |
| pET52b-PA5301-sF | **TTAAGAAGGAGATATACC**ATGGACGTCGGTGCTCGTCT | *PA5301* overexpression |
| pET52b-PA5301-sR | **CTACCGCGTGGCACCAGAGCGAG**GAAATTTGCGGGCGTGGTGG | *PA5301* overexpression |
| pEXG2-mut-PA0534-BS-sF1 | **GGTCGACTCTAGAGGATCCCC**GTTCGACCTCATCTGCCAGG | EMSA (P*PA0534* mut BS) |
| pEXG2-mut-PA0534-BS-sR1 | **GGTTATTTGAGCTCAAAAATAA**CACTTAAAATTTCACGACACCAG | EMSA (P*PA0534* mut BS) |
| pEXG2-mut-PA0534-BS-sF2 | **TTATTTTT<u>GAGCTC</u>AAATAACC**AATATAATTTCACGGAGGTCGCG | EMSA (P*PA0534* mut BS) |
| pEXG2-mut-PA0534-BS-sR2 | **ACCGAATTC<u>GAGCTC</u>GAGCCC**ACGAAGCGTAGCCCCTCGTA | EMSA (P*PA0534* mut BS) |
| CTX-PA2776-lacZ-sF | **GATATCGAATTCCTGCAGCCC**GGGAAGCGTGAAGAAGTCCG | EMSA (P*PA2776* mut BS) |
| CTX-PA2776-mut-lacZ-sR | **GCTAGTTAGTTAGGATCCCCC**TGCGGCATCTTGCACCTC<u>GCA</u>TTTTT <u>GC</u>TTTTT<u>CAC</u> GATGCCCGGATGCTACCCGA | EMSA (P*PA2776* mut BS) |
| CTX-mdpA-lacZ-sF | **GATATCGAATTCCTGCAGCCC**CCGCATTGATCAGGTTGCCG | EMSA (P*mdpA* mut BS) |
| CTX-mdpA-mut-lacZ-sR | **GCTAGTTAGTTAGGATCCCCC**TGTTCTCCTCGTCTGGACCGGGCTGA AATTTAAGTCGAG<u>AACTG</u>ATTCAGGTCAAAACAAATTTTCAGT | EMSA (P*mdpA* mut BS) |
| (Cy5-) pergAB-EMSA-F | (Cy5-) CAGCCTTCTCCCGATGGCAGT | EMSA (Trouillon *et al.*, 2020) |
| pPA0534-EMSA-F | CAGCCTTCTCCCGATGGCAGT CTCGGGACTGGTGTCGTGAA | EMSA |
| pPA0534-EMSA-R | CAGCCTTCTCCCGATGGCAGT AGGAACATCGCGACCTCCGT | EMSA |
| pPA1360-EMSA-F | CAGCCTTCTCCCGATGGCAGT GACATGTTCGAGGACGCTGG | EMSA |
| pPA1360-EMSA-R | CAGCCTTCTCCCGATGGCAGT CGGGAAACGGGATGTGCACT | EMSA |

| | | |
|---|---|---|
| pPA1885-EMSA-F | CAGCCTTCTCCCGATGGCAGT TATCCATGAGTGGCCCTGGC | EMSA |
| pPA1885-EMSA-R | CAGCCTTCTCCCGATGGCAGT GGGCAAGCTCCAATGCAAGG | EMSA |
| pPA2776-EMSA-F | CAGCCTTCTCCCGATGGCAGT CGATTTCGACTCGGGTAGCAT | EMSA |
| pPA2776-EMSA-2-R | CAGCCTTCTCCCGATGGCAGT TGCGGCATCTTGCACCTCTG | EMSA |
| pPA4498-EMSA-F | CAGCCTTCTCCCGATGGCAGT CCCGAACCCCTATTTTTAAGTG | EMSA |
| pPA4498-EMSA-R | CAGCCTTCTCCCGATGGCAGT TGTTCTCCTCGTCTGGACCG | EMSA |
| pPA4985-6-EMSA-F | CAGCCTTCTCCCGATGGCAGT ACGGGACAGAAACTATCCAGTC | EMSA |
| pPA4985-6-EMSA-R | CAGCCTTCTCCCGATGGCAGT GCGCTTCAGAAACAGGATATGA | EMSA |
| uvrD up | CATATCCTGGTGGACGAGTTCC | RT-qPCR |
| uvrD down | CGCTGAACTGCTGGATGTTCTC | RT-qPCR |
| qPCR_pauB1_up | ACCTATGGACGGCGATCCTG | RT-qPCR |
| qPCR_pauB1_down | ACGAAGCGTAGCCCCTCGTA | RT-qPCR |
| qPCR_fumC1_up2 | TCGGGCAACTTCGAACTGAA | RT-qPCR |
| qPCR_fumC1_down2 | GAGCTTGCCCTGGTTGACCT | RT-qPCR |
| qPCR_PA0265_up2 | TGTTCCGCTTCAAGGACGAG | RT-qPCR |
| qPCR_PA0265_down2 | CCATGCCGTACTCCAGTTGC | RT-qPCR |
| qPCR_PA1360_up | TCCCTGGCGAAAAGCATGT | RT-qPCR |
| qPCR_PA1360_down | AGGAGCAGCAGCAGGATCAG | RT-qPCR |
| qPCR_PA1541_up | TCGCCTACGCCCTTTGGGAA | RT-qPCR |
| qPCR_PA1541_down | TACAGGCCGATGCTTTCGCC | RT-qPCR |
| qPCR_PA1885_up2 | CCGAACGGCTGTACCTGAAG | RT-qPCR |
| qPCR_PA1885_down2 | CAGCCAGGCGCTTGTAGAAA | RT-qPCR |
| qPCR_PA4498_up2 | GCATCAGCACCACCGAAGTC | RT-qPCR |
| qPCR_PA4498_down2 | ACCGAACAGGACGATGCAGA | RT-qPCR |
| qPCR_PA4500_up | CAACGCCGACGATGTGCTGT | RT-qPCR |
| qPCR_PA4500_down | TTGTCCAGGCCCATGTCGGT | RT-qPCR |
| qPCR_PA4985_up | CGTGGTGATGACGTCCACCT | RT-qPCR |
| qPCR_PA4985_down | ACCACCGCCCAGTAGTCCAT | RT-qPCR |
| qPCR_PA4986_up | CTGTTCAGCCCCCTGGAAAT | RT-qPCR |
| qPCR_PA4986_down | GTGGCTCGTTGACCAGGTTG | RT-qPCR |
